# Live access to the emotional dynamics of REM sleep dreams in lucid dreamers with narcolepsy

**DOI:** 10.1101/2025.08.27.672557

**Authors:** Jean-Baptiste Maranci, Pierre Champetier, Delphine Oudiette, Emma Chabani, Luc Masset, Alix Romier, Augustin Chen, Amelie Barbier, Umberto Pannacci, Federica Sapo, Başak Türker, Smaranda Leu-Semenescu, Andrea Pinna, Isabelle Arnulf

## Abstract

Sleep helps regulate emotions, but it is still unclear whether -and how- the emotions we experience in dreams contribute to this regulation. To uncover the potential function of dream emotions, we must first understand what they are and how they unfold in dreams. The emotional content of dreams has mostly been studied using post-sleep dream reports, which provide a biased and static snapshot of a complex and dynamic experience. In this study, we took a more direct approach, accessing dream emotions in real-time. We asked twenty-four lucid dreamers with narcolepsy to report the emotional valence of their dreams, - positive, negative or neutral-, while still asleep, using predefined facial codes during daytime naps monitored with polysomnography. Of the 126 naps recorded, 62 contained at least one emotional code during REM sleep, yielding 191 codes in total. The ratios of positive and negative codes were evenly balanced per nap. The 33 naps with at least two codes allowed us to track the dream emotional dynamics. Over half of these naps showed opposite emotional valences (positive and negative). By measuring the time elapsed between codes, we estimated the average duration of a given dream’s emotional valence in REM sleep to be about one minute. Positive emotions emerged on average earlier than negative ones during lucid REM sleep. These findings confirm the highly emotional nature of dreams and, more importantly, highlight that emotions in REM sleep dreams are fluid and fast-changing. Such emotional dynamics during REM sleep dreams may help us to better understand the mechanisms of the emotional regulatory function of dreams.

## Introduction

What do we dream about and why? The nature of dreams is enigmatic, and the majority of them remain elusive to us. To ascertain whether they serve a function, it is first necessary to gain a clear understanding of their content. A variety of methodologies have been utilized to examine dream content, including last dream recall, general dream questionnaires, dream diaries, and provoked awakening in sleep labs, with the latter being regarded as the gold standard.^1^ Notably, research highlights the prevalence of emotions in dreams, suggesting that they may be a crucial element.^2^ This emotional aspect has shaped several theories aimed at explaining the function of dreams. Some theories suggest that emotions experienced during dreams assist in processing emotional memories formed during wakefulness^3–8^ or prepare individuals for future threatening situations.^9^ If these theories are validated, elucidating the mechanisms by which emotions are processed during dreams could have significant implications for understanding some psychiatric disorders characterized by dysregulated emotional processing, such as depression. Yet, the emotional landscape of dreams remains mysterious and warrants further investigation.

A seminal study of 1,000 dream reports collected between 1947 and 1950 evaluated by external judges found a predominance of negative emotions.^10^ Subsequent studies have nuanced this predominance of negative emotions. Notably, studies comparing dream emotions scored by external judges with the subject’s self-assessment have shown that external rating tend to underestimate dream emotions, especially positive ones.^11–13^ The self-evaluation of dreams obtained through awakenings during rapid eye movement (REM) sleep has shown a preponderance of positive emotions.^11,14^ In contrast, self-evaluations in dream diaries have shown either a balanced ratio of positive and negative emotions,^15,16^ a predominance of positive emotions,^13,14^ or a predominance of negative emotions.^17^ The differences could be partly explained by the impact of the location (hospital versus home) or the type of scale used.

A common limitation of all these approaches is that they rely on the subject’s report obtained after awakening. Such reports are subject to potential biases, such as amnesia and the reconstruction of dream content.^1^ Furthermore, it should be noted that reports of dream emotions present a static and simplified representation of dreams, omitting the complexity and dynamic aspects of dreaming. A few studies have explored the sequence of emotions in a dream in a more detailed way, asking participants to write in a dream diary each dream scene with the corresponding emotions.^17^ Their findings revealed that 46% of dreams containing at least one emotion featured a “major emotional shift,” defined as a transition from a positive to a negative emotion or vice versa. Additionally, the authors noted that negative emotions were more prevalent towards the end of the reports. Zadra et al. analysed reports of bad dreams (defined as disturbing, non-arousing dream) and found that 38% of bad dreams ended positively.^18^ Nightmares (defined as disturbing, arousing dreams) were less likely to end positively (22% of the time). These studies suggest that a notable proportion of dreams encompass a combination of positive and negative emotions that alternate over time. However, crucial limitations of these approaches include memory bias and the absence of temporal measurement, which impedes the ability to assess the frequency of emotional shifts and the duration of emotional states within a dream.

To address these limitations, our team studied the emotional behaviour of people with REM sleep behaviour disorder (RBD), a condition characterized by dream enactment due to the partial destruction of the REM sleep paralysis system. Negative and positive behaviours (visible on the face of the participants) were observed in a balanced way during REM sleep.^19^ Consistent with reports from healthy subjects, a third of the REM sleep episodes with emotional behaviours exhibited both positive and negative emotional behaviours.^20^ However, contrary to what healthy dream reports suggest, negative behaviours were observed earlier in REM sleep episodes compared to positive (and neutral) behaviours. This new approach has limitations too, as the exact correspondence between emotional behaviour and subjectively felt emotion during dreams remains to be explored. Additionally, emotional behaviours are relatively rare in RBD patients, representing a limited window into REM sleep emotions.

An alternative approach to accessing the subjective component of a dream in real time is lucid dreaming. A lucid dream is defined as a dream in which the dreamer is aware of dreaming. In some cases, it is also possible for the dreamer to communicate information from the dream using pre-defined codes made with the eyes or by contracting the muscles of the face.^21–23^ The chances of obtaining such codes from a lucid dream are highest in participants with narcolepsy,^24^ a disease whose cardinal symptom is irrepressible daytime sleepiness, leading to naps that often include REM sleep.^25^

In the present study, we employed lucid dreaming in participants with narcolepsy to examine the emotional dynamics of dreams. This approach provided an avenue to access emotions subjectively experienced in a dream state. We recruited 24 participants with narcolepsy who reported experiencing lucid dreams several times per week. They were instructed to signal the emotion experienced in the ongoing lucid dream, using pre-defined muscle codes during daytime naps.

## Results

### Lucid dreamers with narcolepsy can successfully report the emotional valence of their ongoing dreams

The objective of this study was to access the emotional valence and dynamics of ongoing dreams in real-time. We recruited 24 participants diagnosed with narcolepsy (14 females; mean age 30 ± 10 years), all of whom reported experiencing several lucid dreams per week. During daytime naps, participants were instructed to indicate the emotional valence of their ongoing dreams through specific muscle contractions: three consecutive frowns for a negative dream content, three smiles for a positive dream content, and alternating between a frown and a smile for a neutral dream content **(Figure 1)**. They were monitored using polysomnography (PSG), including electroencephalography, electrooculography, electromyography, respiratory plethysmography, and electrocardiography. Isometric muscle contractions were recorded using two EMG channels placed on the right zygomaticus (for smiling) muscle and the corrugator muscle (for frowning), as described in previous studies.^22,23^ Participants were then awakened from their naps either after a predefined duration or at REM sleep termination in 2 different recording sessions. During the first session, 3 naps per participant were recorded, each with a predefined duration of 30 minutes. These naps were preceded and followed by the presentation of images selected from the *International Affective Picture System* (IAPS),^26^ consisting of 40 negative and 40 neutral images for each block (80 images in total). In the second session, the number of naps varied, and the maximum nap duration was reduced to 20 minutes, without the visualization of IAPS images (see Methods for details). After awaking, participants were interviewed about 1) the content of their dreams, 2) whether they were lucid at any point, 3) the codes they produced, if any, and 3) the specific order in which they produced these codes and the corresponding dream content, if remembered.

**Figure 1.**
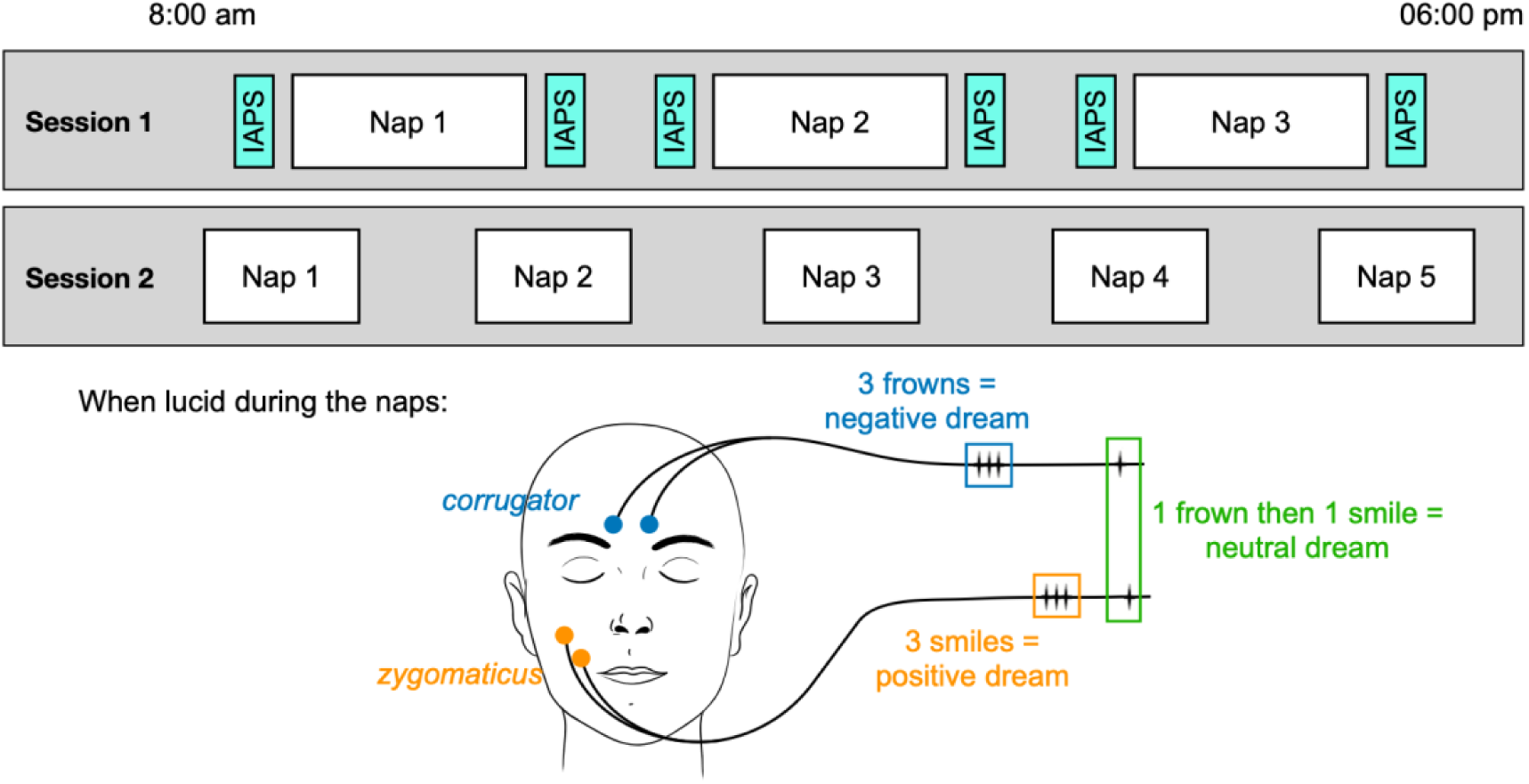
Experimental design of the study. Participants were instructed to take daytime naps and report the emotional valence of their ongoing dreams if they became lucid. The predefined codes were as follows: 3 frowns for a negative dream content (blue), 3 smiles for a positive dream content (orange), and alternating a frown then a smile for a neutral dream content (green). Two different sessions of recording were conducted. In Session 1, three 30-minute naps were recorded, each preceded and followed by visualization of images from the *International Affective Picture System* (IAPS). In Session 2, up to five naps of 20 minutes maximum were recorded over one or two successive days.

We analysed a total of 126 naps (mean ± SD = 5.3 ± 4.1 per participant; median = 3, IQR = [3 – 5]; range = [2 - 13]). The codes were scored by visual inspection of the corrugator and zygomatic EMG by two independent judges, who were blinded to the participant identity, sleep stage, and post-nap report. To be conservative, only codes selected by both judges were considered for further analysis. A total of 191 codes were selected during REM sleep (79 negative, 56 positive, and 56 neutral). Additionally, we observed 77 codes during periods of short arousals or awakenings (27 positive, 18 negative, and 32 neutral), and 7 codes during NREM sleep, with 3 occurring in stage N1 (2 negative and 1 neutral) and 4 in stage N2 (3 positive and 1 neutral). Only REM sleep codes were considered for further analysis. When considering all the naps, 62/126 (49%) presented at least one code during REM sleep, with an average of 1.5 ± 3 codes per nap (range [0 – 23]). Naps with at least one REM sleep code were followed by a report of lucidity after awakening in 88.7% of the cases (N = 55/62), and participants reported having realized a code in 79% of the cases (N = 49/62). A more detailed description of the correspondence between codes and their reports after awakening can be found in **Figure S1.**

At the participant level, 71% of participants (17/24) produced at least one code during REM sleep, with a median of 5, IQR = [3 – 9] codes per participant who produced codes (mean 11 ± 17; range = [1 – 69]). Notably, two participants achieved an exceptional performance, generating 69 REM sleep codes (36.1% of total REM sleep codes) and 40 REM sleep codes (20.9% of total REM sleep codes), respectively. These two participants were qualified as “champion lucid dreamers” (ChLD) 1 and 2. **Data S, Table 1** provides additional details on the naps and codes during REM sleep for each participant. **Figure 2** presents examples of the correspondence between observed codes in REM sleep and the corresponding dream report obtained after the end of the nap, an additional example can be found in **Figure S2**.

**Figure 2.**
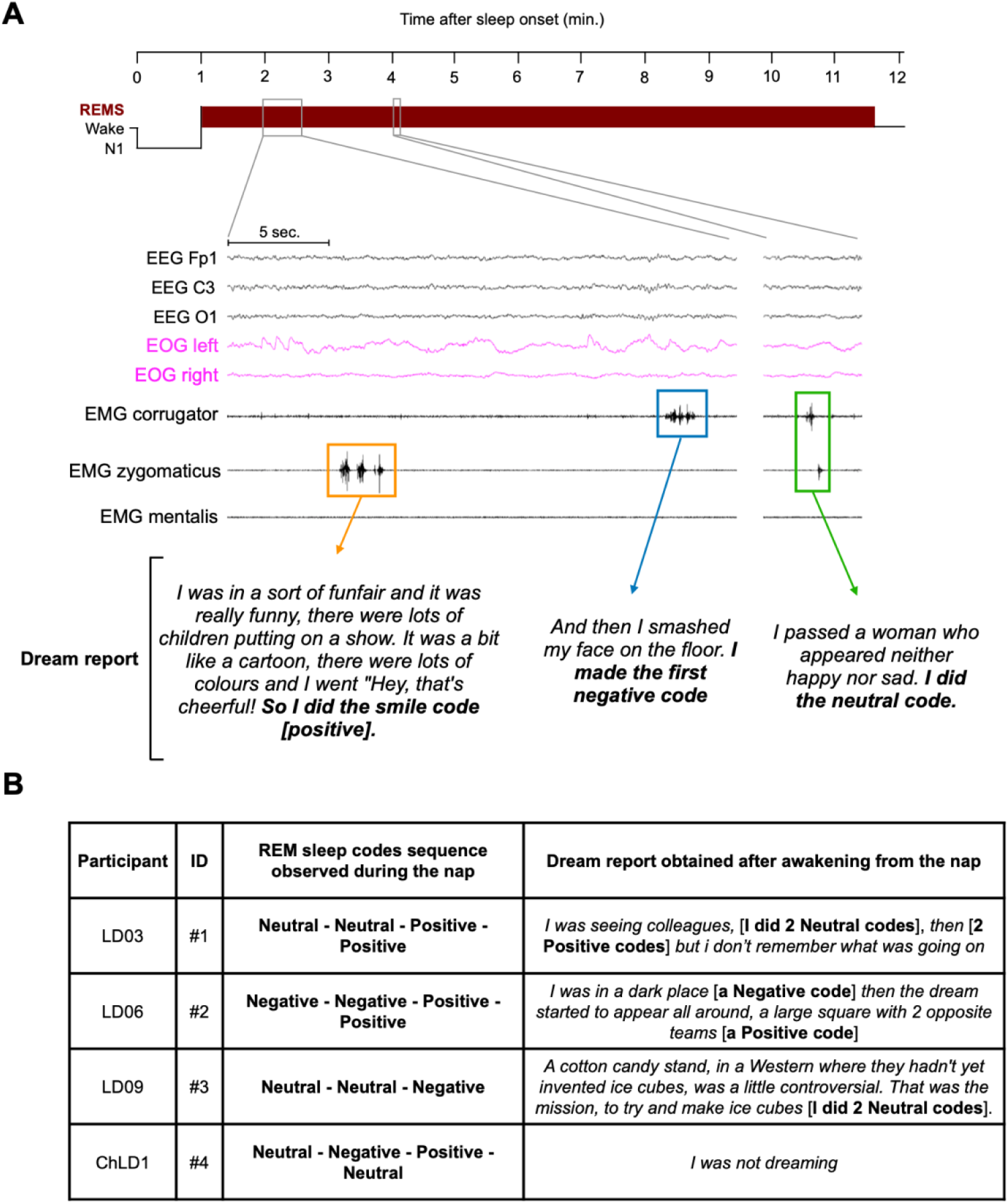
Narcoleptic lucid dreamers can successfully report the emotional valence of their ongoing dreams. **(A)** An illustrative example of a nap with a REM sleep onset typical of a participant with narcolepsy, the extracts illustrate the succession of a positive code (orange), a negative code (blue), at some distance a neutral code (green), with the corresponding dream report. **(B)** Illustrative examples of REM sleep code sequences with the corresponding dream reports, which show either a perfect correspondence (ID#1), a correct sequence of reported code valence but a mismatch for the number of codes realised (ID#2), a match for neutral codes but a negative code not reported (ID#3) and a complete absence of dream report following a nap with observed codes (ID#4). REMS = REM sleep.

### Balanced emotional ratio and mixed emotions in lucid REM sleep

We initially sought to ascertain whether REM sleep dreams are as emotionally charged as previous research on dream reports has indicated. Additionally, we aimed to determine the proportion of positive and negative emotions present in such dreams. For each nap with codes (N = 62), a code ratio for a given valence was calculated by dividing the number of codes of a given valence during the nap by the total number of codes observed, all valence combined. When using the nap as the unit of analysis, the ratio of negative codes (49 ± 42%) was higher than the ratio of neutral codes (15 ± 27%, Wilcoxon test: W = 2812, p < 0.0001) and tended to be higher than the ratio of positive codes (36 ± 40%, Wilcoxon test: W = 2245.5, p = 0.09). Additionally, the ratio of positive codes was higher than the ratio of neutral codes (Wilcoxon test: W = 1290, p = 0.0009), **(Figure 3A)**. Given that some naps were preceded by the visualization of negative and neutral IAPS images (N = 27/62, 44%) and others were not (N = 35/62, 56%), we aimed to test whether this exposure influenced the valence ratio of the codes during the naps. To achieve this, we conducted a mixed linear model with the ratio of a given valence as the dependent variable and the presence versus absence of IAPS image visualization as the independent variable. The result indicated a significant effect of IAPS image visualisation with an increased negative ratio after watching the IAPS images (61 ± 40% with IAPS images versus 39 ± 42% without, df = 52.8, t score = 2, p = 0.049). The model did not converge for the positive ratio and a simple t test was used with a trend found showing a decreased positive ratio after watching the IAPS images (26 ± 38% with IAPS images versus 44 ± 40% without, df = 57.8, t score = 1.82, p = 0.07) and no difference was found for neutral ratio (13 ± 25% with IAPS images versus 17 ± 29% without, df = 53.3, t score = -0.3.34, p = 0.64). Overall, this indicates that image visualization may influence the emotional ratio of naps, with an increased negative ratio. Therefore, we compared the ratios of different valences only for naps without image visualization. This time, the negative and positive valence ratios did not differ and were higher than the neutral valence ratio (**Figure 3B**). To ensure that the absence of a difference between negative and positive valence ratios was not due to insufficient statistical power, we conducted a Bayesian mixed model with the ratio as the dependent variable, valence (positive versus negative) as a fixed regressor, and nap identity as a random factor showing a result in favour of the null hypothesis (evidence of no difference, i.e., BF01 > 3) (BF01 = 3.7).

**Figure 3.**
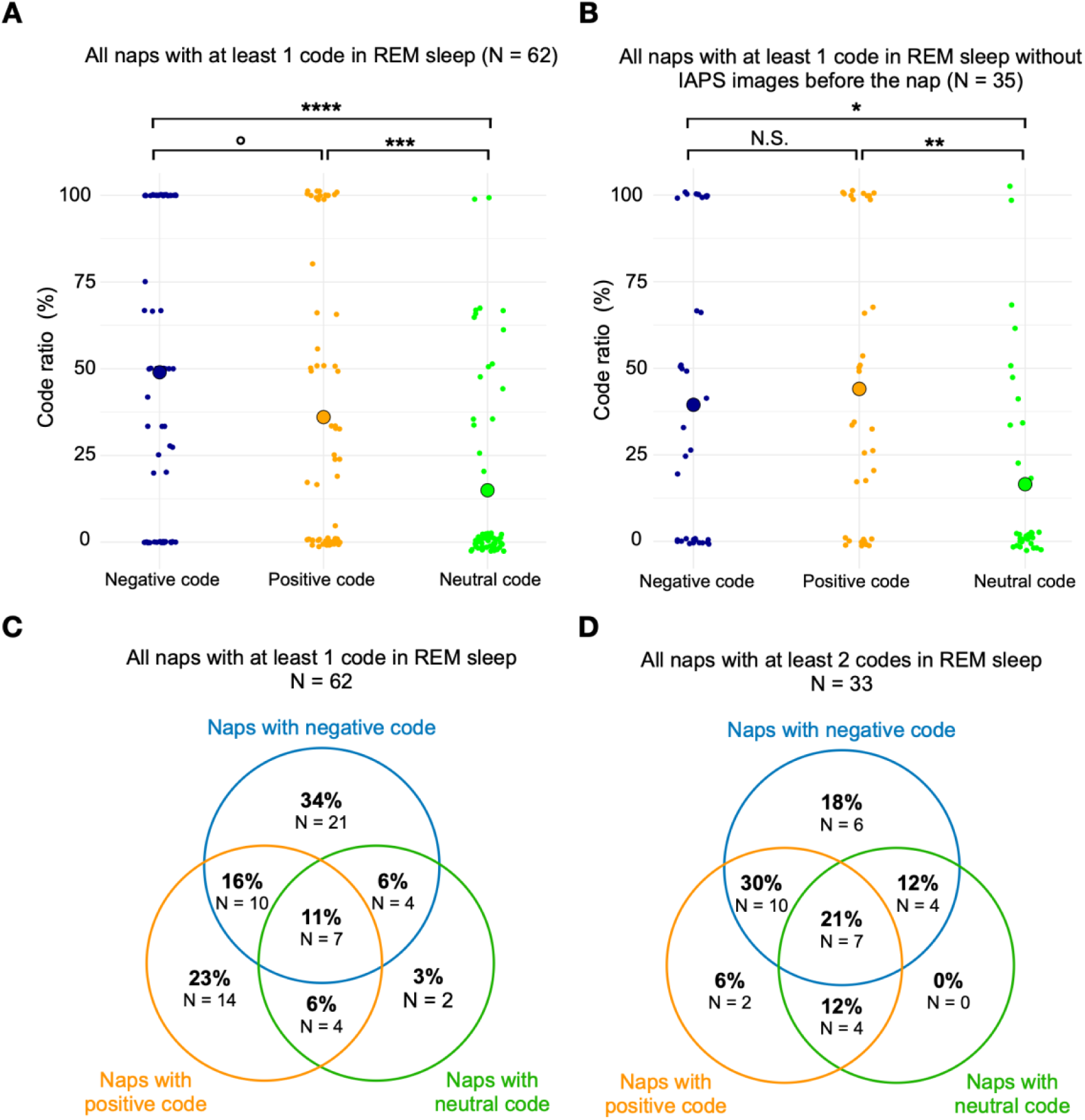
Balanced emotional ratio and mixed emotions in lucid REM sleep. **(A)** Distribution of the ratio of a given REM sleep code valence within a nap, calculated by dividing the number of REM sleep codes of a given valence by the total number of REM sleep codes within a nap. Each dot represents a nap, with larger dots representing the mean ratio for a given valence among the naps. **(B)** Distribution of the ratio of a given REM sleep code valence within a nap limited to the naps not preceded by IAPS images before the nap. **(C)** Representation of the number and percentage of naps with at least one code of a given valence, among all the naps in which at least one REM sleep code observed. Values outside the circle’s intersections indicate that only codes of a given valence were observed in REM sleep. Values at the intersection of two circles indicate that two different code valences were observed in REM sleep during the nap. Values in the centre, at the intersection of the three circles, indicate that all the different codes were observed in REM sleep **(D)**. Representation of the number and percentage of naps with at least one code of a given valence, among all the naps in which at least two REM sleep code observed. IAPS = *International Affective Picture System;* °p < 0.1; ***p < 0.001; ****p < 0.0001.

We then analyzed whether different valence codes could be identified within the same REM sleep nap and at what frequency. The number and proportion of naps with REM sleep codes of a given valence or combinations of different valence codes are presented in **Figure 3C**. When considering all naps containing REM sleep codes (N= 62), 39% presented at least two codes of different valence. Among the 33 naps with two or more REM sleep codes (i.e., the minimum condition to observe two codes of different valence within the same nap), 76% exhibited two different code valences, including codes of opposite valence (negative/positive) in 52% of the naps **(Figure 3D)**.

### The shift in emotional valence occurs within a relatively brief time frame

To explore the dynamics of valence changes in more detail, we analysed the time intervals between two consecutive REM sleep codes within the same nap. We distinguished between inter-code intervals where two successive codes had the same valence, defined as “valence continuity” and those with differing valences, defined as “valence shift.” Shifts between two opposite valences (negative/positive) were classified as “opposite valence shift,” and shifts from a negative or a positive code to a neutral code or from a neutral code to a positive or a negative code were classified as “non-opposite valence shift”. Among the participants, 12/24 (50%) had more than one muscular code in REM sleep during a given nap. A total of 130 inter-code intervals were observed, 108 of which occurred within a continuous REM sleep sequence (i.e., without any arousals or awakenings between two successive codes). Of these, 72/130 (55%) exhibited valence continuity (55%), while 58/130 (45%) demonstrated a valence shift. Of the latter, 22 were opposite valence shifts (17%). An example of two successive opposite valence shifts is show in **Figure 4**. All different combinations of inter-code intervals are detailed in **Figure 5A**.

**Figure 4.**
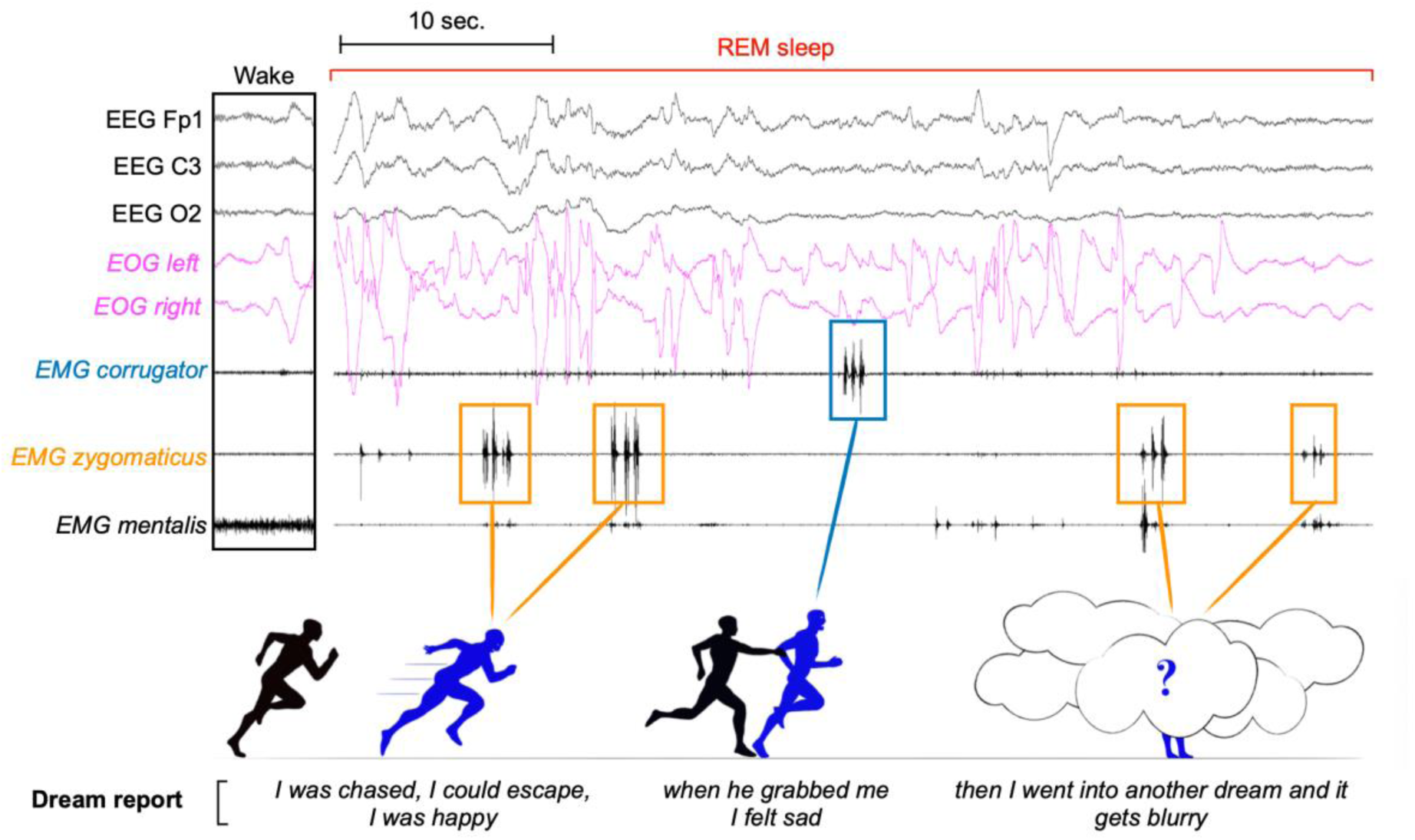
Example of two successive opposite emotional shift in lucid REM sleep. The left side of the figure depicts a wake period from the same participant for comparative purposes. The participant is in REM sleep and produces two successive positive codes, followed by a negative code a few seconds later, and then two successive positive codes. The dream report content associated by the participant with the codes after awakening from the nap is indicated. It should be noted that the content for the last two codes could not be detailed as the participant only recalls a change in dream.

**Figure 5.**
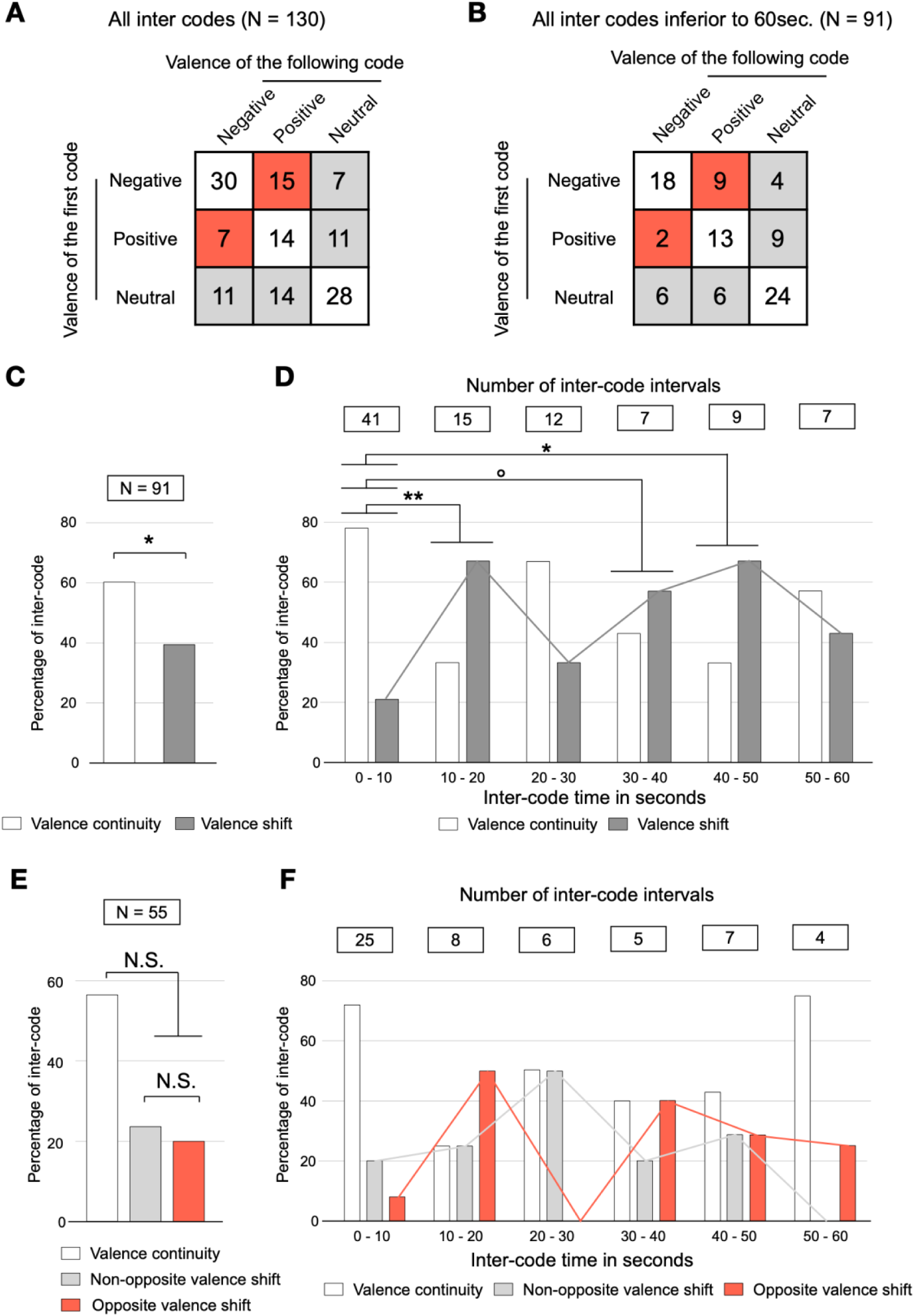
Valence shift in a short time scale. **(A)** All combinations of all inter-code observations considering the valence of the first and the following REM sleep code. **(B)** All combinations of inter-code intervals with a duration of less than 60 seconds, considering the valence of the first and the following REM sleep code. **(C)** Percentages of inter-code intervals (≤ 60 seconds) with valence continuity and valence shift. **(D)** Percentages of inter-code intervals with valence continuity and valence shift in successive 10-second time windows, considering the time elapsed between two successive codes. **(E)** Percentages of inter-code intervals (≤ 60 seconds) with a positive or negative initial code, showing valence continuity, non-opposite valence shift, and opposite valence shift. **(F)** Percentages of inter-code intervals with a positive or negative initial code, showing valence continuity and valence shift in successive 10-second time windows, considering the time elapsed between two successive codes. N.S.: non-significant;°p < 0.1; *p < 0.05; **p < 0.01.

To analyse how frequently a valence change might occur in REM sleep, we selected inter-codes intervals lasting less than 60 seconds (70% of all inter-code intervals, N=91/130). Among them, 98% (N=89/91) were observed in a non-interrupted REM sleep sequence. All the different combinations of inter-codes observed in less than 60 seconds between two successive codes are indicated in **Figure 5B**. The ratio of continuity (60%) was higher than the ratio of shifts (40%) in a generalized mixed model (estimate valence continuity vs. shift = 0.42, z score = 1.98, p = 0.048) (**Figure 5C**). This result suggests a certain stability of emotional valence over a period of 1 minute, though with a notable rate of valence shifts. To examine if this stability varied with the time intervals between codes, we divided the inter-code intervals into six groups based on their duration, each falling within successive 10-second windows between 0 and 60 seconds. The percentages of valence continuity and valence shifts for each interval are shown in **Figure 5D**. A time window effect was found in a generalized mixed model with valence shift vs. valence continuity as the dependent measure and time window as a fixed regressor (χ²(5) = 12.73, p = 0.03). The percentage of valence continuity (78%) versus shift (22%) was higher in the [0 - 10] sec time window (generalized linear mixed model: estimate continuity vs. shifts = 1.2, z score = 2.43, p = 0.02). No differences in continuity versus shift percentages were found in other time windows (for details see **Data S Table 2**). To test whether the stability of emotional valence (assessed via the percentage of continuity) varied in function of the time elapsed between codes, we contrasted the different time windows with the presence of a shift versus continuity as the dependent variable and the time windows in fixed contrast in a generalized mixed model. All the contrasts are presented in **Data S Table 3**. The percentage of shifts was higher in the [10-20], [30-40] (with only a trend for the latter), and [40-50] second windows compared with the [0-10] second window **(Figure 5D)**. A trend for a higher percentage of shift was also found for the [10-20] contrasted with the [20-30] time window. In summary, these results suggest that valence remains mostly stable within 10 seconds after a code but becomes less stable in some of the time windows beyond 10 seconds.

We then focused on the inter-code intervals with either an initial negative or positive code to specifically study the dynamics of positive and negative dreams. By focusing on inter-code intervals shorter than 60 seconds, 55 intervals were selected with positive or negative code as the initial code. The percentage of valence shift (44%) did not differ from valence continuity (56%) (generalized mixed model: estimate continuity vs. shift = 0.26, z score = 0.94, p = 0.35) (**Figure 5E**). The percentages of opposite (20%) and non-opposite shifts (24%) did not differ (generalized linear mixed model: estimate opposite vs. non opposite shift = -0.17, z score = -0.41, p = 0.68). As described before, we inspected the ratio of shifts, (non-opposite and opposite shifts) in successive 10 sec time windows, see **Figure 5F**. A time window effect was not found for shift vs. continuity probability (χ²(5) = 6.8, p = 0.24).

Since two ChLD accounted for half of the codes, we ran the same analyses excluding them. A detailed analysis can be found in the **Supplementary Results** section. In summary, valence shifts, notably opposite valence shifts, could be observed. Considering the inter-code intervals lasting less than 60 seconds, the higher ratio of continuity versus shift observed in previous analyses was not reproduced. There were too few codes to perform a detailed statistical analysis for each 10-second interval.

### One minute of emotional valence stability in lucid REM Sleep

We aimed to estimate the duration of emotional valence during REM sleep. We considered only uninterrupted REM sleep sequences (without awakenings or short arousals) with at least two successive codes. When two consecutive codes had the same emotional valence, we checked whether a subsequent code followed and ignored intermediate codes as long as they presented the same valence, until reaching: 1) a code with a different valence, at which point the elapsed time was measured and classified as the end of the current valence (shift); or: 2) an interruption of REM sleep, at which point the time between the two most distant codes of the same valence was measured and classified as an unachieved valence end (no shift observed). Using this approach, 64 sequences were defined and measured, with 44 showing a shift and 22 showing no shift before an interruption of REM sleep.

These measures were used to run a survival analysis of the valence. The median survival times for all valences combined and for each valence considered separately are presented in **Figure 6A**, while the corresponding survival curves are shown in **Figure 6B**. The median survival time was 51 sec when considering all valences together. A Cox mixed model showed a trend for an effect of the valence of the code on survival probability (Chi2 = 5.27, df = 2, p = 0.07). The probability of a shift tended to be higher for a positive versus negative valence (Hazard ratio positive vs. negative = 2.12, z score = 1.85, p = 0.07) but was not different for neutral vs. negative (Hazard ratio neutral vs. negative = 1.16, z score = 0.38, p = 0.7) or for positive vs. neutral (Hazard ratio positive vs. neutral = 1.83, z score = 1.49, p = 0.14).

**Figure 6.**
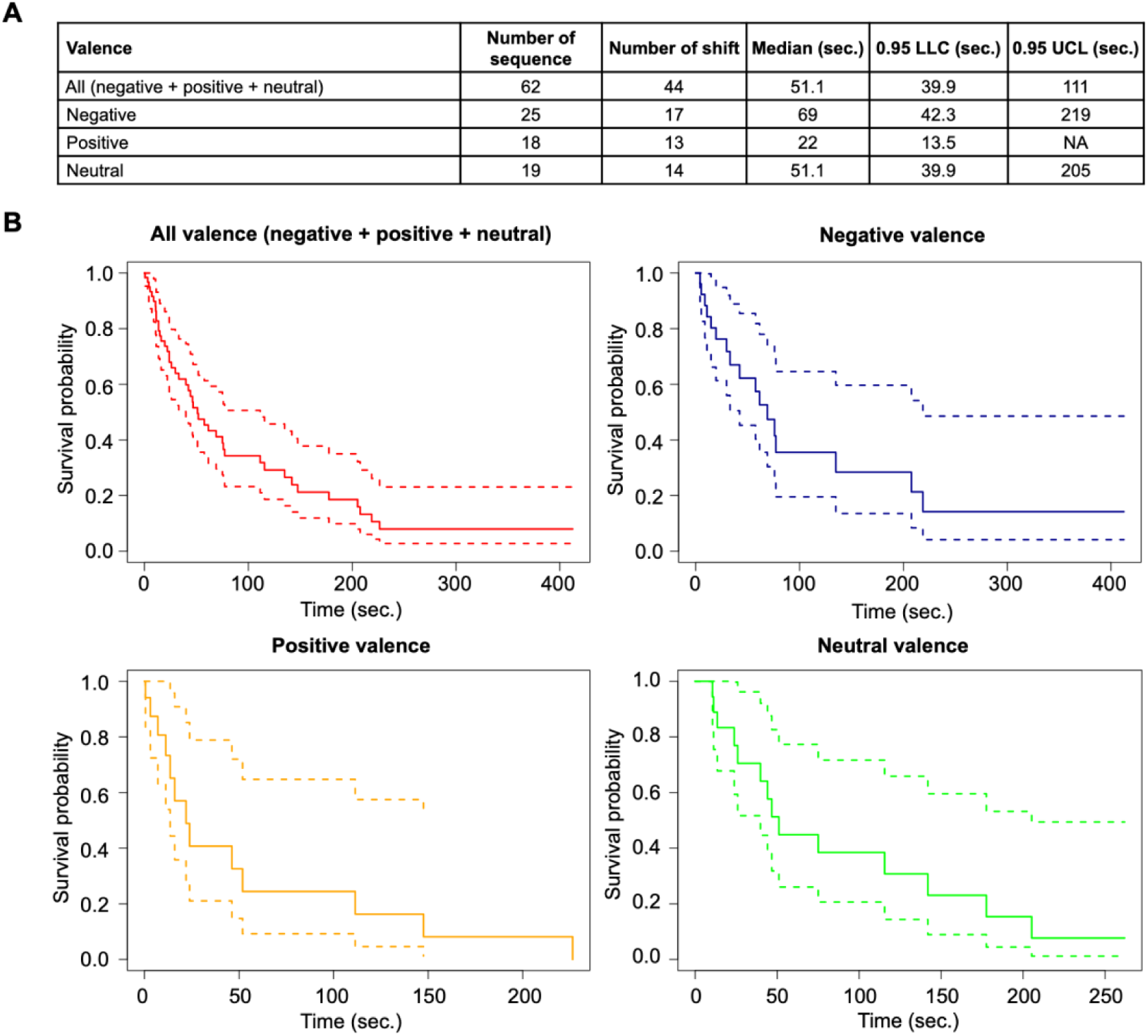
One minute of valence stability in lucid REM Sleep. **(A)** Table showing, from left to right, the number of sequences analysed for survival analysis, the number of sequences with valence shift, the median time calculated for the survival model (in sec), the 95% and the lower confidence limit (LCL) and the 95% upper confidence limit (UCL), for all valences combined and then for each given valence (negative, positive and neutral). **(B)** Corresponding survival curves, all valences combined (red), then for negative (blue), positive (orange) and neutral valence (green).

After excluding the two ChLDs from the analysis, 20 sequences were measured with 10 shifts observed and with a median time of 142 seconds in the survival analysis, further details are presented in **Figure S4**. A Cox mixed model did not show an effect of valence on shift probability (Chi2 = 0.47, df = 2, p = 0.79).

### Positive codes precede negative codes in lucid REM sleep

Finally, we wondered whether certain valences occurred relatively earlier or later during REM sleep. For each code, we considered the time elapsed from the start of REM sleep. The mean time of appearance of codes in REM sleep was longer for negative (median = 3.9 min [2.5 min – 7.5 min]; mean 5.9 ± 4.7 min) than positive codes (median = 3.3 min [1.9 min – 5.4 min]; mean 4.2 ± 3.2 min) (generalized linear mixed model: estimate negative vs. positive = 0.28, z score = 2.08, p = 0.04) but not than neutral codes (median = 4.3 min [2.2 min – 6.6 min], mean 4.5 ± 2.8 min) (generalized linear mixed model: estimate negative vs. neutral = 0.15, z score = 1, p = 0.3). No difference in time of appearance in REM sleep was found between positive and neutral codes (generalized linear mixed model: estimate positive vs. neutral = -0.13, z score = -0.87, p = 0.39) (**Figure 7**).

**Figure 7.**
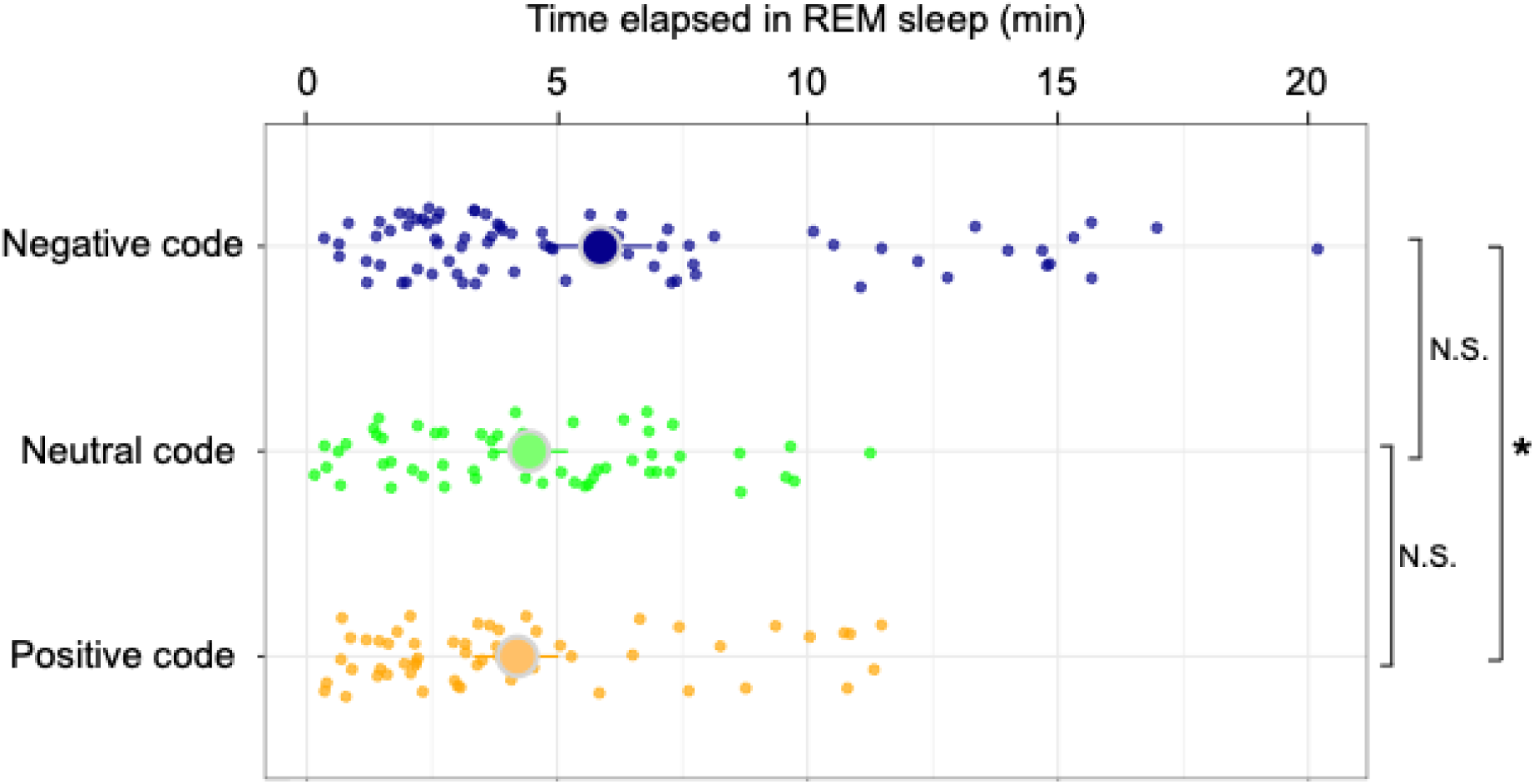
Positive codes precede negative codes in lucid REM sleep. Each point represents a code conspiring its valence (negative, neutral, or positive) with the corresponding time elapsed since the beginning of the REM sleep episode during the naps. The larger point represents the mean and the line the range of the confidence limit. N.S.: non-significant, * p < 0.05.

These results were reproduced after excluding the 2 ChLDs from the analysis, with a longer time of appearance for negative (median = 5.6 min [2.9 min – 8.6 min]; mean 6.5 ± 5 min) versus positive codes (median = 2.2 min [1.5 min – 4.1 min]; mean 3.1 ± 2.4 min) (generalized linear mixed model: estimate negative vs. positive = 0.7, z score = 3.58, p = 0.0003) and a trend versus neutral codes (median = 4.7 min [1.7 min – 6.1 min]; mean 4.5 ± 3.3 min) (generalized linear mixed model: estimate negative vs. neutral = 0.498, z score = 1.781, p = 0.07). No difference was found for the time of appearance of neutral and positive codes (generalized linear mixed model: estimate positive vs. neutral = -0.21, z score = -0.682, p = 0.495) (**Figure S5**).

## Discussion

In this study, we accessed the emotional content of ongoing lucid dreams in participants with narcolepsy, using pre-defined facial codes. Most dreams were reported as emotional (versus neutral) with a balanced ratio of positive and negative codes. Most naps (with at least 2 codes observed in REM sleep) showed different or even opposite valences. Dream emotions showed rapid changes, and a valence stability could be estimated to be about one minute. Positive emotions occurred earlier than negative emotions in REM sleep.

We observed a trend towards a higher ratio of negative versus positive emotions, but this effect could be explained by the viewing of negative (and neutral) images before some naps. Naps without previously viewed images showed no difference in the ratio between positive and negative emotion. In any case, the ratio of both negative and positive emotions was higher than the ratio of neutral valence. Our results confirm the idea that dreams are highly emotional and that the high rates of emotional dreams reported after awakenings from REM sleep are not due to memory biases.^11,14^ A balanced ratio of positive and negative emotions was found in some dream diary studies.^12,15,16^ However, our results contrast with two studies that found a higher rate of positive emotions in dreams with awakenings induced after 5 or 10 minutes of REM sleep.^11,14^ Compared to these studies, our measurements are not biased by the limitations of dream reports, such as amnesia and reconstruction. However, our study model could present a bias towards negative emotion (see the limitations section).

Our results extend the understanding of the dynamics of emotions in dreams. We observed transitions between different valences over short temporal durations during sleep. Limiting our analysis to inter-code intervals shorter than one minute, the valence change rate was about 40%, and more specifically for emotional valence (not neutral) the transition to an opposite valence was about 20%. These findings are consistent with the observation of Merritt et al.,^17^ who observed transitions from one emotion to its opposite (negative to positive or vice versa) in dream diaries annotated with the corresponding emotions experienced. Our approach introduces several new insights. Firstly, in addition to the emotions (positive/negative), we investigated the neutral aspect of dreams, making the study of emotional dynamics more complete. In the approach of Merritt et al., the neutrality of a dream segment was not scored, making it difficult to determine whether the described change in opposite emotional valence was separated by a neutral segment or not. We found a balanced ratio of transition to neutral and opposite shift (24% and 20% respectively). Crucially, our approach allowed us to measure the objective times elapsed between valences and to ensure that the transitions described occurred within a short time interval, specifically less than one minute in the analysed measures.

Our findings indicate that the average duration of an emotional valence during a dream is approximately one minute, with an increase to two minutes when excluding two participants. To our knowledge, no previous study based on dream reports has ever asked to evaluate the duration of an emotion experienced during a dream. Therefore, this provides the first insight into this question. We lacked a contrast to quantify how much emotion fluctuates in REM sleep compared to wakefulness. It is challenging to conceive a method to measure the emotional state of awake participants with such frequency and ecological validity, as the participants would have to report their emotional state several times a minute. In the waking state, emotions can occur very briefly, this is one of their characteristics (distinguishing them from mood, which is a prolonged state), however, such a frequent occurrence of emotions, sometimes opposite, seems intuitively improbable. The reason for such brevity of emotions during dreams was not investigated in our study. However, it can be hypothesized that this characteristic might be important for the emotional regulation function attributed to dreams.^3–8^ Processing negative emotions may require alternating between reactivating negative emotions and engaging with neutral or positive experiences. This process could avoid a “negative overload” that could overwhelm our emotional regulation capacities and potentially trigger a nightmare. One element in favour of this hypothesis is the study carried out by Zadra et al. showing that bad dreams more often have a positive ending than awakening nightmares.^18^ This suggests that positive emotions experienced during dreams may serve as a protective mechanism that prevents bad dreams from transforming into nightmares.

We found negative codes to appear later than positive codes, although their appearance windows overlapped, especially at the beginning of REM sleep. This result is consistent with the study of dream narratives by Merritt et al., showing a predominance of negative content at the end of dream narratives.^17^ This result is also consistent with the findings of Fosse et al., who observed a higher ratio of negative emotions following provoked awakenings in the second half of REM sleep episodes compared to the first half during daytime naps in patients with narcolepsy.^27^ However, this result contrasts with our previous study of emotional behaviours in patients with RBD, where negative emotional behaviours occurred earlier in REM sleep.^20^ This discrepancy may be explained by the behavioural nature of the emotions studied in RBD patients, which may not completely align with the subjective emotions of dreams, or by specific changes inherent to the RBD model, which is prodromal to degenerative diseases. This result could suggest that an excess of negative dreams might occur when REM sleep episodes are prolonged, as observed in patients suffering from depression.^28^ Based on our previous hypothesis, a relatively high ratio of negative emotions after an extended period of REM sleep could represent a “critical moment” for the formation of nightmares. Investigating the timing of nightmares during a REM sleep episode versus an awakening with a simple dream reported in a home recording paradigm, as conducted by Paul et al.,^29^ could help explore this hypothesis.

The main limitation of our study is that our results may not be applicable to other contexts. First, the participants had narcolepsy, and their dreams were studied during daytime naps. A comparison of dream emotions in patients with narcolepsy, self-rated after provoked awakenings during REM sleep in daytime naps, showed that the prevalence of fear/anxiety was twice as high as in dreams collected during night-time REM sleep awakenings in healthy subjects.^27^ However, there were no major differences in the other emotions studied (negative or positive). The dream emotions in our study could therefore be partially biased towards the negative compared to the dreams of healthy subjects. Second, we focused here on lucid dreams, which could also bias the type of emotion obtained, since some studies show that lucid dreams are more positive than non-lucid dreams.^30,31^ The control over the content of the dream that lucidity sometimes allows could also influence the emotion obtained, but we asked the participants not to influence the content of the dream in progress and only to report its valence through the codes. Third, the number of NREM sleep codes in our study was insufficient to perform analyses on NREM sleep valence ratio and dynamics. The distribution of codes was found to be highly uneven, with two participants accounting for half of the analyzed codes. However, all analyses using the codes (or inter-code intervals) were re-run excluding these two participants.

Our study paves the way for future research on emotions in dreams. Firstly, it serves as a proof of concept that dream emotions can be studied in real-time using lucid dreaming. Our findings could be replicated in healthy, non-narcoleptic subjects trained in lucidity in a sleep laboratory. Paradigms aiming to alter the emotional content of dreams through odors or sounds could be tested, with results evaluated directly from an ongoing dream. Previous paradigms investigating the EEG correlates of dream emotions based on dream reports could also benefit from this approach, directly adjacent to codes of a given valence for optimal temporal resolution.^32,33^ Our research demonstrates the dynamic nature of dream valence, encouraging further study of this aspect of dreams. A crucial aspect would be to investigate this dynamic in patients suffering from mental disorders such as depression, nightmares, or post-traumatic stress disorder, compared to healthy subjects. Thus, our study opens the door to the investigation of the emotional dynamics of dreams, which remains underexplored.

## Methodology

### Participants

Twenty-four participants with narcolepsy were recruited for the study (14 females; mean age: 30.3 ± 10.4 years). All were frequent lucid dreamers with several lucid dreams reported per week. Of the participants, 18 had narcolepsy type 1 (with cataplexy or hypocretin deficiency) and 6 had narcolepsy type 2 (without cataplexy or hypocretin deficiency). All were patients diagnosed in the National Reference Center for Narcolepsy in the Pitié-Salpêtrière university Hospital according the international criteria for narcolepsy including: 1) excessive daytime sleepiness occurring daily for at least 3 months; 2) a mean sleep latency ≤ 8 min and two or more sleep onset REM sleep periods on the multiple sleep latency tests (five tests performed at 08:00, 10:00, 12:00, 14:00 and 16:00; and 3) no other better cause for these findings, including sleep apnoea syndrome, insufficient sleep, delayed sleep phase disorder, depression and the effect of medication or substances or their withdrawal. The participants were instructed to discontinue their treatments the day prior to the experiment.

### Experimental Design

The recordings of the participants were conducted in two sessions at the sleep unit of Pitié Salpêtrière University Hospital in Paris. During Session 1, 22 participants underwent three 30-minute daytime naps each. Prior to and following each nap, participants viewed and rated 40 negative and 40 neutral images from the IAPS (results not reported here). One participant did not complete the IAPS task and had four naps, while another had an additional 10-minute nap in a session without IAPS images. Technical issues prevented nap analysis for two participants, resulting in 66 naps analysed for Session 1.

In Session 2, 7 participants (five of whom participated in Session 1) were recorded over one or two consecutive days based on availability, with four to five naps per day. Sleep was monitored in real time. If a code was observed during REM sleep, participants were awakened if they showed signs of exiting REM sleep (cessation of eye movements for more than one minute or appearance of spindles or K-complexes). If no code was observed, participants were awakened after a 20-minute nap. This approach was used to maximise the likelihood of capturing memories of codes performed and associated dream content. One participant completed four naps in a single day, while others completed nine or ten naps over two days. A total of 60 naps were recorded for Session 2.

Prior to each nap in both sessions, participants were instructed to report the emotional valence of their ongoing dream if lucid: three smiles for positive dreams, three frowns for negative dreams, and alternating smiles and frowns for neutral dreams. Participants were instructed not to attempt to alter dream content. Subsequently, upon awakening from naps, participants were questioned about: 1) dream content; 2) instances of lucidity; 3) any codes produced; and 4) the sequence of code generation relative to dream content. During Session 2, they were also asked to evaluate the time elapsed between two reported successive codes, in seconds.

### Sleep scoring

Sleep stages were scored offline by a certified sleep expert according to established guidelines, using Profusion® software (Compumedics®, Medical Data Technology, Australia). For scoring, the EEG and EOG signals were filtered between 0.3 and 15 Hz, the EMG and EKG signals were filtered between 10–100 Hz and 0.3–70 Hz respectively. A 50 Hz notch filter was applied on all channels. Sleep scoring was visually performed on 30-s time epochs, each scored as wakefulness, N1, N2, N3 or REM sleep, according to the American Academy of Sleep Medicine international rules.

### Code selection

Two independent judges examined the EMG channels of the corrugator and zygomatic muscles in 30-second epochs, tasked with selecting codes. They were blinded to participant identity, sleep stage, and post-nap reports. Both judges reviewed all recording epochs. Their agreement rate for codes identified during REM sleep was 86%. As a conservative measure, any code lacking consensus between the judges was excluded from the analysis.

### Time elapsed between codes measures

When two REM sleep codes occurred during the same nap, the time elapsed between the end of the first code and the start of the next code was measured in seconds. It was noted whether the valence of the two codes was the same and termed “valence continuity” or different “valence shift” and in this case whether the valences of the codes were opposite (positive to negative or negative to positive) termed “opposite valence shift”.

A different measurement was used for survival analyses. When two successive codes showed a continuity of valence, we checked for the presence of a following code, ignoring the successive codes as long as they presented the same valence, until reaching: 1) a code with a different valence, at which point the elapsed time was measured and classified as the end of the current valence (shift); or 2) an interruption of REM sleep, at which point the time between the two most distant codes of the same valence was measured and classified as an unachieved valence end (no shift observed).

### Statistical analysis

All statistical analyses were conducted using R and its various packages detailed below.^34^ As most of the quantitative measures analysed did not follow a normal distribution and occasionally exhibited extreme values, we presented most measures using both median and quartiles, as well as mean and standard deviation. Comparison of valence ratios during naps was conducted using a simple Wilcoxon test, with naps as the unit of analysis. The effect of IAPS image visualisation was assessed using the valence ratio as a dependant measure, image visualisation as a fixed effect regressor and nap identity as random effect with the ‘lme4’ package.^35^ Analysis of the non-normally distributed quantitative measures was performed using a generalized linear mixed-effects model with the ‘glmmTMB’ package,^36^ which included a log transformation of the measures. The unit of analysis was the code and/or the inter-code interval, with the subject’s identity as the random factor in the model. Evaluation of valence duration was estimated using survival analysis to calculate the median duration with the ‘survival’ package.^37^ Survival comparison among different valences was analysed using a mixed-effects survival model with the ‘coxme’ package.^38^ Occasionally, Bayesian statistics were used to prove the null hypothesis with the calculation of the Bayes factor 01 (BF01), which indicates a level of evidence for the null hypothesis when it is greater than 3. The ‘BayesFactor’ package was then used.^39^ All analyses using code or inter-code interval as the unit of analysis were systematically rerun if possible, excluding the two participants with the highest number of codes (ChLD 1 and 2).

## Acknowledgements

The promotor of the study was ADOREPS (Association pour le développement et l’organisation de la recherche en pneumologie et sur le sommeil). This study was funded by the Agence Nationale de la Recherche (ANR-20-CE37-0001-01 grant to D.O.) and a research grant from Société Française de Recherche et Médecine du Sommeil (to D.O. and E.C.). J.B.M. received a grant from Assistance Publique - Hôpitaux de Paris (APHP) and Sorbonne University (‘Poste d’accueil APHP’). The National Reference Center for Narcolepsy benefits from a recurrent grant from the National Program on Rare Diseases (PNMR-3 grant to I.A.) from the Health Ministry, which encourages research on narcolepsy. The research leading to these results has received funding from the national program “Investissements d’avenir” ANR-10-IAIHU-0006.

## Author contributions

J.B.M., P.C., E.C., A.R., A.B. and B.T. collected the data. J.B.M, U.P, F.S, L.M., conducted the data analysis. A.C. participated to the conception of the study. A.P., I.A., and D.O. provided supervision and guidance throughout the study. J.B.M., I.A., A.P., and D.O. wrote the manuscript.

## Supplemental Results

### The shift in emotional valence occurs within a relatively brief time frame

Since two ChLD accounted for half of the codes, we ran the same analyses exposed in the “**The shift in emotional valence occurs within a relatively brief time frame”** section, excluding them. A total of 39 inter-code intervals were observed (including 29 in a non-interrupted REM sleep sequence), 54% with valence continuity (N=17/39) and 66% of valence shift (N=22/39) and 26% of opposite shift (N=10/39). Details of the different combinations of codes are shown in **Figure S3A.** Among them, 21 were shorter than 60 seconds (**Figure S3B**). The valence shift ratio (35%) did not differ from the continuity ratio (65%) (generalised linear mixed model: estimate shift vs. continuity = -0.69, z score = -1.5, p = 0.13) (**Figure S3C**). The number of inter-code intervals observed in successive 10-second time windows was too low for statistical analysis. Therefore, the percentages of shifts and continuities observed are presented only descriptively in **Figure S3D**.

Excluding the 2 ChLD, 32 inter-codes with an initial negative or positive code were observed, of which 17 (53%) lasted less than 60 seconds. The percentages of valence shift (35%) and valence continuity (65%) did not differ (generalized linear model: estimate shift versus continuity = -0.6061, z score = -1.194, p = 0.23) (**Figure 3E**). We did not compare the percentage of opposite valence shift versus the percentage of non-opposite valence shift due to the low number of occurrences. The percentages according to the 10 sec time windows are shown only descriptively in **Figure S3F**.

**Data S Table 1.**
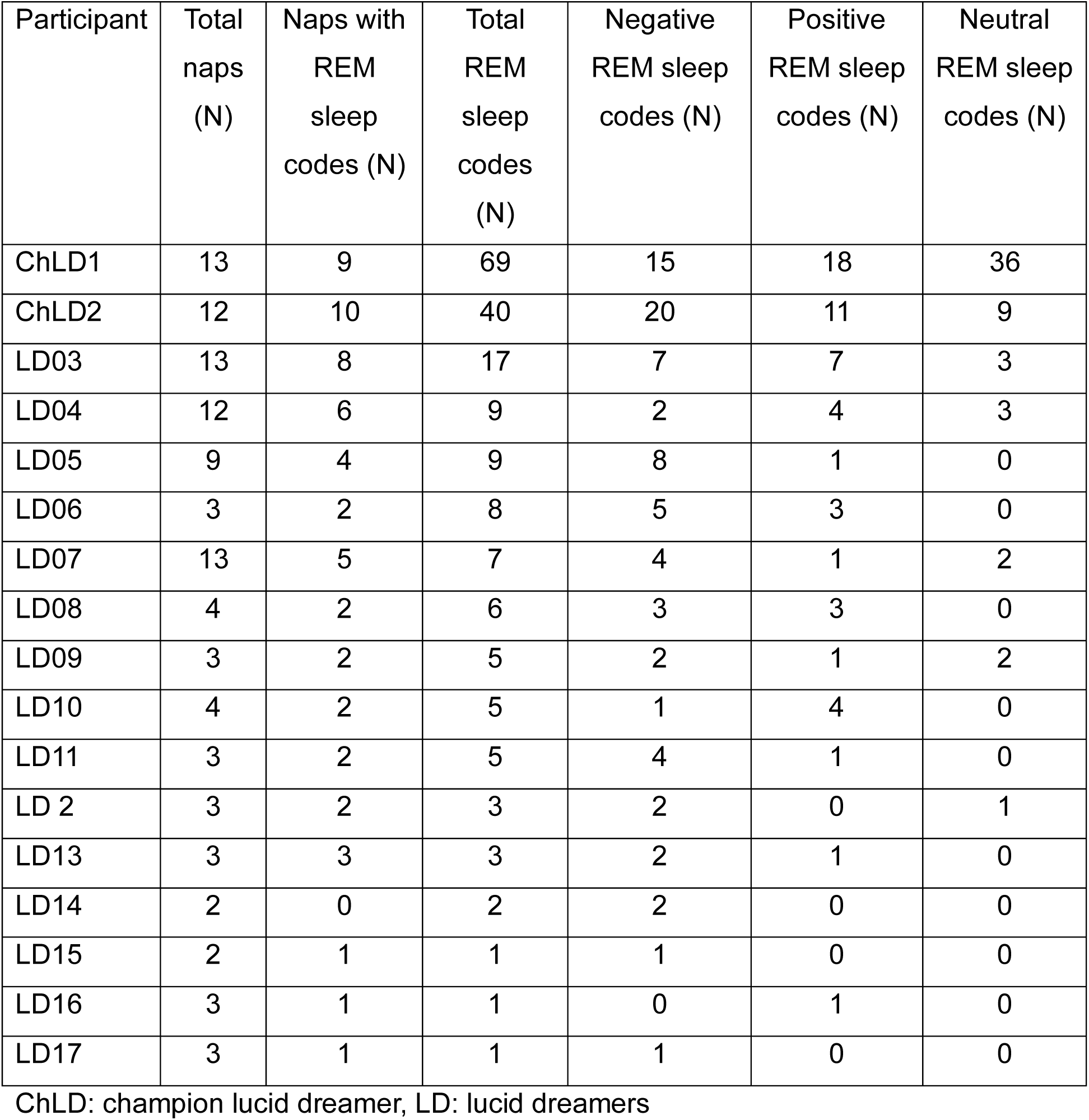
List of participants who performed at least one code during REM sleep, with the total number of naps analysed, the number of naps with REM sleep codes, the total number of codes selected, and the number of codes of a given emotional valence (negative, positive or neutral).

| Participant | Total naps (N) | Naps with REM sleep codes (N) | Total REM sleep codes (N) | Negative REM sleep codes (N) | Positive REM sleep codes (N) | Neutral REM sleep codes (N) |
| --- | --- | --- | --- | --- | --- | --- |
| ChLD1 | 13 | 9 | 69 | 15 | 18 | 36 |
| ChLD2 | 12 | 10 | 40 | 20 | 11 | 9 |
| LD03 | 13 | 8 | 17 | 7 | 7 | 3 |
| LD04 | 12 | 6 | 9 | 2 | 4 | 3 |
| LD05 | 9 | 4 | 9 | 8 | 1 | 0 |
| LD06 | 3 | 2 | 8 | 5 | 3 | 0 |
| LD07 | 13 | 5 | 7 | 4 | 1 | 2 |
| LD08 | 4 | 2 | 6 | 3 | 3 | 0 |
| LD09 | 3 | 2 | 5 | 2 | 1 | 2 |
| LD10 | 4 | 2 | 5 | 1 | 4 | 0 |
| LD11 | 3 | 2 | 5 | 4 | 1 | 0 |
| LD 2 | 3 | 2 | 3 | 2 | 0 | 1 |
| LD13 | 3 | 3 | 3 | 2 | 1 | 0 |
| LD14 | 2 | 0 | 2 | 2 | 0 | 0 |
| LD15 | 2 | 1 | 1 | 1 | 0 | 0 |
| LD16 | 3 | 1 | 1 | 0 | 1 | 0 |
| LD17 | 3 | 1 | 1 | 1 | 0 | 0 |

ChLD: champion lucid dreamer, LD: lucid dreamers

**Data S Table 2.** Models contrasting the percentage of valence continuity versus valence shift observed in successive 10-second time windows, considering the distance between two successive codes separated by less than 60 seconds.

| Model | Estimate continuity<br>versus shift | Z score | P value |
| --- | --- | --- | --- |
| Shift in [0-10] time<br>window ~ 1 +<br>(1\|Participant) | 1.2 | 2.43 | **0.02** |
| Shift in [10-20] time<br>window ~ 1 +<br>(1\|Participant) | -0.69 | -1.27 | 0.21 |
| Shift in [20-30] time<br>window ~ 1 +<br>(1\|Participant) | -0.69 | 1.13 | 0.26 |
| Shift in [30-40] time<br>window ~ 1 +<br>(1\|Participant) | -0.29 | 0.38 | 0.71 |
| Shift in [40-50] time<br>window ~ 1 +<br>(1\|Participant) | -0.69 | 0.98 | 0.33 |
| Shift in [50-60] time<br>window ~ 1 +<br>(1\|Participant) | 0.29 | 0.38 | 0.71 |

Bold characters indicate a p value < 0.05.

**Data S Table 3.** Contrast between different 10 seconds time windows characterising the distance between two successive codes on their effect on the probability of valence shift versus valence continuity.

| Model | Time windows contrast | Estimate for an effect on shift versus continuity | Z score | P value |
| --- | --- | --- | --- | --- |
| Contrast against the [0 - 10] time window |  |  |  |  |
| Shift ~ Time window + (1 \| Participant) | [0 – 10] over [10 – 20] | -1.96 | -2.95 | **0.003** |
| Shift ~ Time window + (1 \| Participant) | [0 – 10] over [20 – 30] | -0.58 | -0.80 | 0.42 |
| Shift ~ Time window + (1 \| Participant) | [0 – 10] over [30 – 40] | -1.56 | -1.83 | 0.07 |
| Shift ~ Time window + (1 \| Participant) | [0 – 10] over [30 – 40] | -1.962 | -2.45 | **0.01** |
| Shift ~ Time window + (1 \| Participant) | [0 – 10] over [40 – 50] | -0.98 | -1.15 | 0.25 |
| Contrast against the [10 - 20] time window |  |  |  |  |
| Shift ~ Time window + (1 \| Participant) | [10 – 20] over [20 – 30] | 1.39 | 1.69 | 0.09 |
| Shift ~ Time window + (1 \| Participant) | [10 – 20] over [30 – 40] | 0.41 | 0.43 | 0.67 |
| Shift ~ Time window + (1 \| Participant) | [10 – 20] over [40 – 50] | 0.00 | 0.00 | 1 |
| Shift ~ Time window + (1 \| Participant) | [10 – 20] over [50 – 60] | 0.981 | 1.04 | 0.3 |
| Contrast against the [20 - 30] time window |  |  |  |  |
| Shift ~ Time window + (1 \| Participant) | [20 – 30] over [30 – 40] | -0.98 | -1 | 0.32 |
| Shift ~ Time window + (1 \| Participant) | [20 – 30] over [40 – 50] | -1.39 | -1.48 | 0.14 |
| Shift ~ Time window + (1 \| Participant) | [20 – 30] over [50 – 60] | -0.41 | -0.41 | 0.68 |
| Contrast against the [30 - 40] time window |  |  |  |  |
| Shift ~ Time window +<br>(1 \| Participant) | [30 – 40] over<br>[40 – 50] | -0.41 | -0.39 | 0.7 |
| Shift ~ Time window +<br>(1 \| Participant) | [30 – 40] over<br>[50 – 60] | 0.58 | 0.53 | 0.59 |
| Contrast against the [40 - 50] time window |  |  |  |  |
| Shift ~ Time window +<br>(1 \| Participant) | [40 – 50] over<br>[50 – 60] | 0.981 | 0.94 | 0.35 |

Bold characters indicate a p value < 0.05.

**Figure S1.**
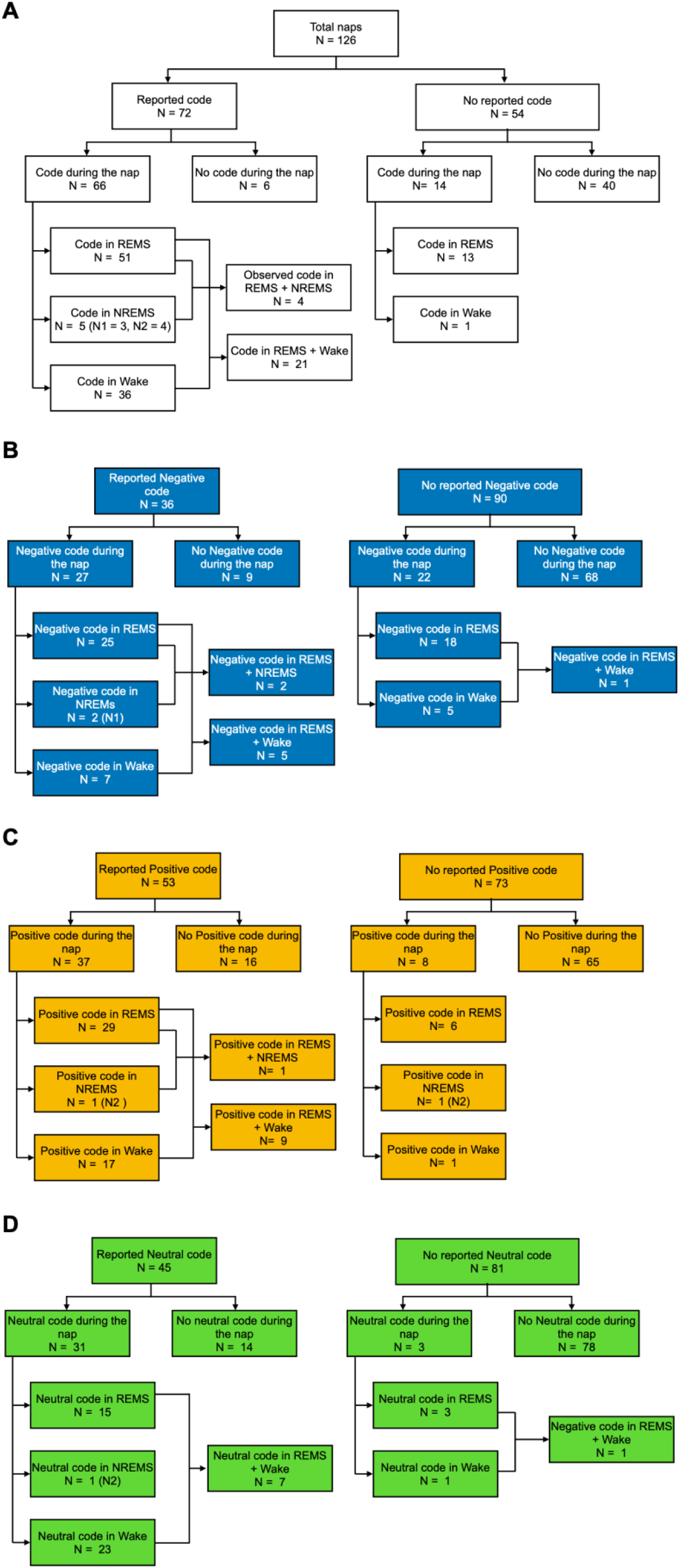
Association between reported codes and code selected during the naps. **(A)** Flowchart detailing the repartition of the naps with or without reported codes after awakening from the nap, further divided depending on whether a code was observed during the nap in REM sleep, NREM sleep or in a wake state (short arousal or a wake epoch) or a combination of these possibilities. **(B)** Flowchart of the repartition of the naps considering only reported or not reported negative codes (blue) further divided into on whether a code was observed during the nap in REM sleep, NREM sleep or in a wake state (short arousal or a wake epoch) or a combination of these possibilities. **(C)** Flowchart of the repartition of the naps considering only reported or not reported positive codes (orange) further divided into on whether a code was observed during the nap in REM sleep, NREM sleep or in a wake state (short arousal or a wake epoch) or a combination of these possibilities. **(D)** Flowchart of the repartition of the naps considering only reported or not reported neutral codes (green) further divided into on whether a code was observed during the nap in REM sleep, NREM sleep or in a wake state (short arousal or a wake epoch) or a combination of these possibilities. NREMS = NREM sleep; REMS = REM sleep.

**Figure S2.**
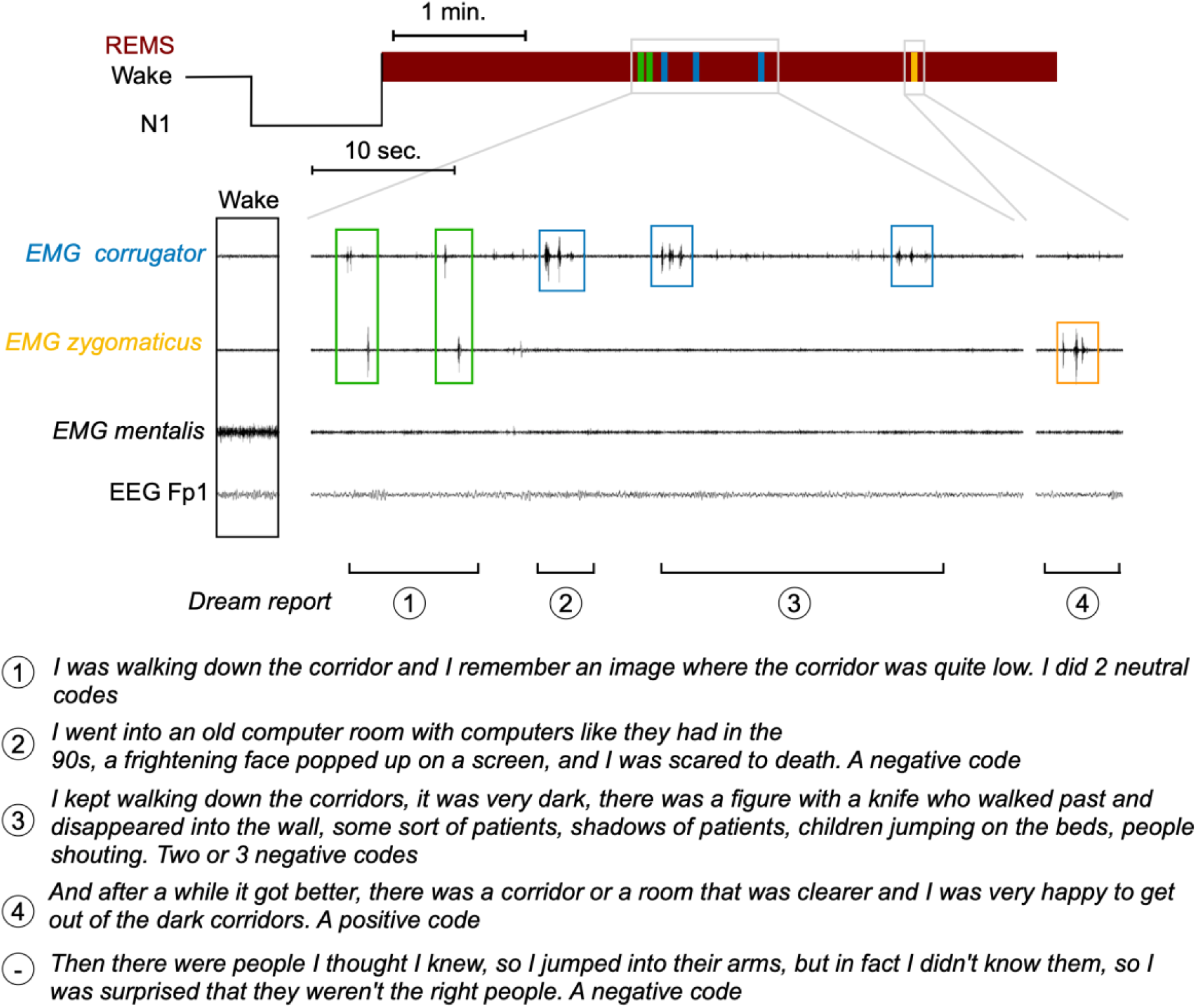
Example of the correspondence between the codes (observed by experimentators or reported by participants after awakening) and associated dream peripeties. The observed sequence of codes (neutral - neutral - negative - negative - negative - positive) partially corresponds to the sequence reported after awakening (neutral - neutral - negative - negative [- negative?] - positive - negative).

**Figure S3.**
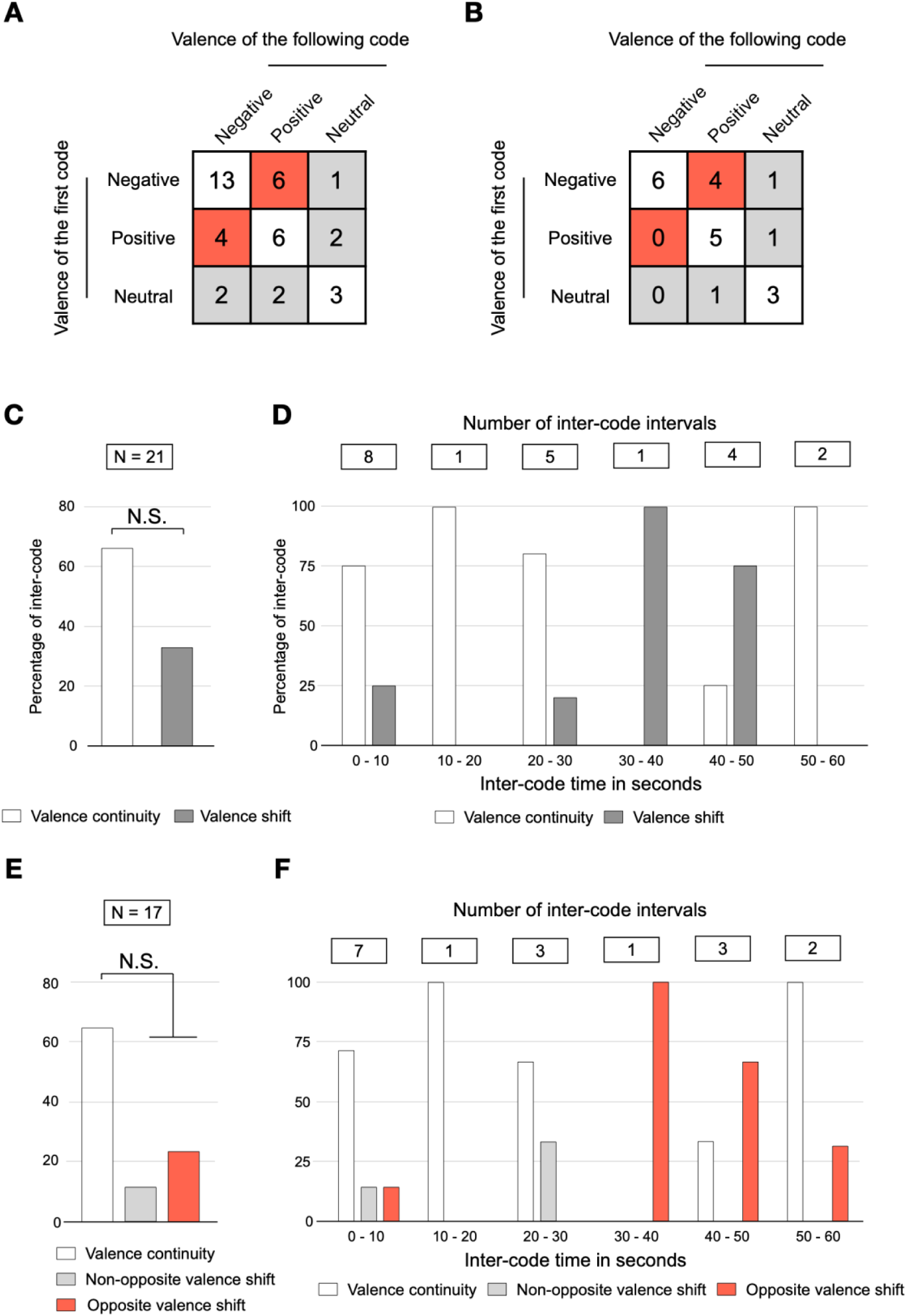
Valence shift in a short time scale without the 2 champion lucid dreamers. **(A)** All combinations of all inter-code observations considering the valence of the first and the following REM sleep code. **(B)** All combinations of inter-code with a duration of less than 60 seconds, considering the valence of the first and the following REM sleep code. **(C)** Percentages of inter-code (shorter than 60 seconds) with valence continuity and valence shift. **(D)** Percentages of inter-code with valence continuity and valence shift in successive 10-second time windows, considering the time elapsed between two successive codes. **(E)** Percentages of inter-code (shorter than 60 seconds) with a positive or negative initial code, showing valence continuity, non-opposite valence shift, and opposite valence shift. **(F)** Percentages of inter-code with a positive or negative initial code, showing valence continuity and valence shift in successive 10-second time windows, considering the time elapsed between two successive codes. N.S.: non-significant; *p < 0.05.

**Figure S4.**
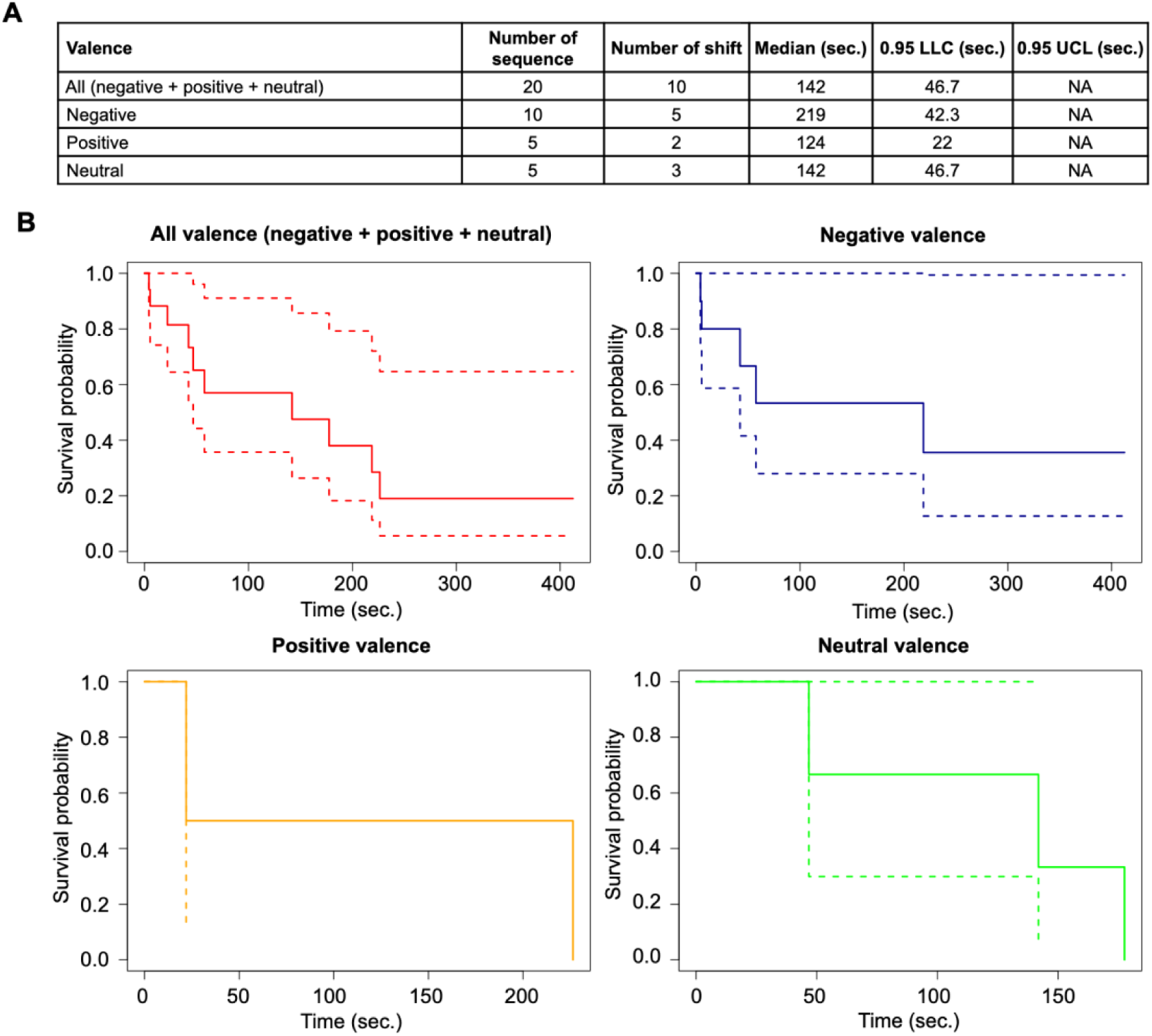
One minute of valence stability in lucid REM sleep. Analysis excluding the 2 champion lucid dreamers. **(A)** Table showing, from left to right, the number of sequences analysed for survival analysis, the number of sequences with an observed valence shift, the median time calculated for the survival model (in sec), the 95% and the lower confidence limit (LCL) and the 95% upper confidence limit (UCL), for all valences combined and then for each given valence (negative, positive and neutral). **(B)** Corresponding survival curves, all valences combined (red), then for negative (blue), positive (orange) and neutral valence (green).

**Figure S5.**
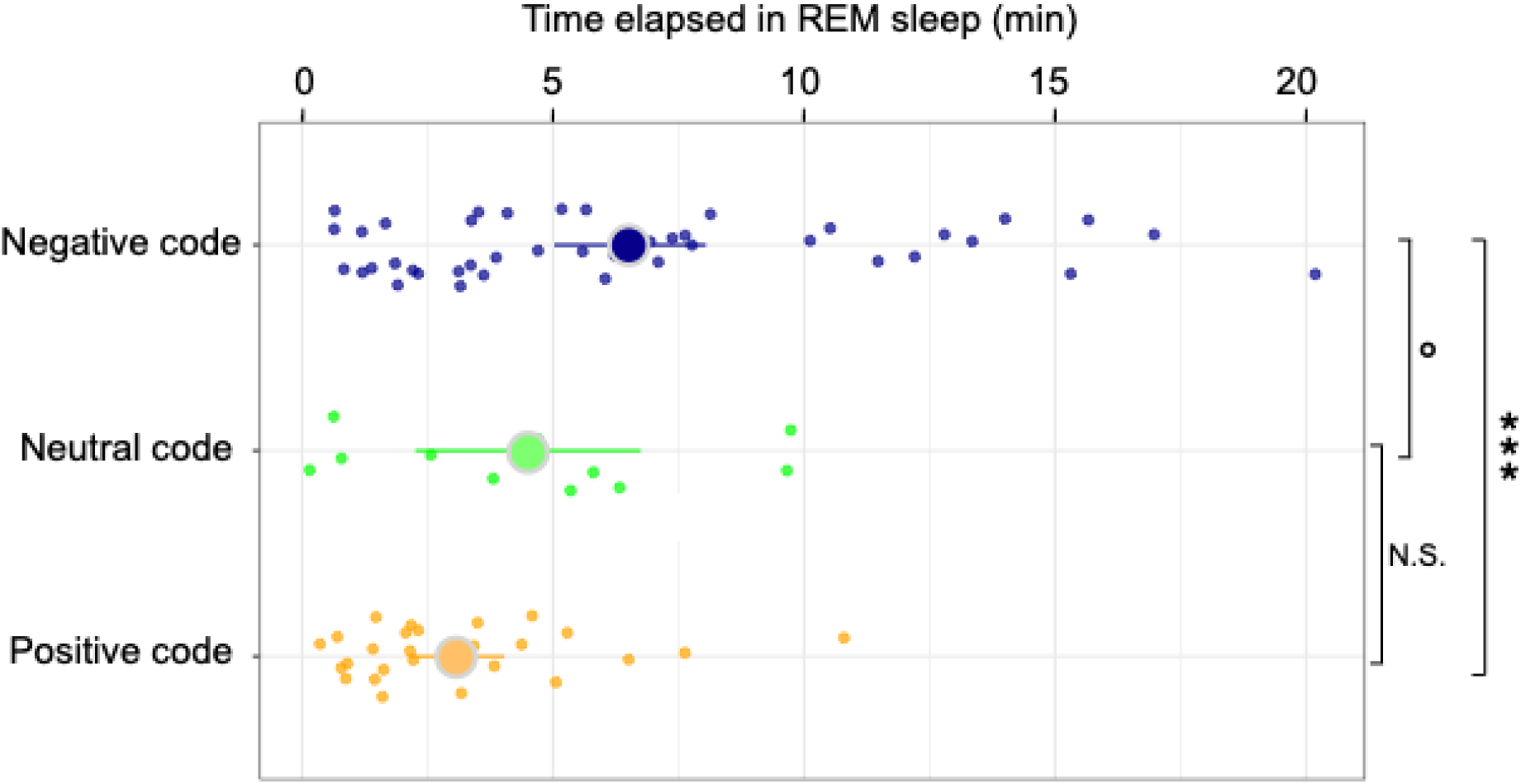
Positive codes precede negative codes in lucid REM sleep. Analysis without the champion lucid dreamers. Each point represents a code conspiring its valence (negative, neutral, or positive) with the corresponding time elapsed since the beginning of the REM sleep episode during the naps. The larger point represents the mean and the line the range of the confidence limit. N.S.: non-significant, °p < 0.05; ***p < 0.001.

